# Genome sequence assembly and annotation of *MATA* and *MATB* strains of *Yarrowia lipolytica*

**DOI:** 10.1101/2025.03.24.645116

**Authors:** Narges Zali, Osama El Demerdash, Kapeel Chougule, Zhenyuan Lu, Doreen Ware, Bruce Stillman

**Affiliations:** Cold Spring Harbor Laboratory, 1 Bungtown Road, Cold Spring Harbor, NY 11724; Graduate Program in Genetics, Stony Brook University, Stony Brook, NY 11794; USDA-ARS Robert W. Holley Center for Agriculture and Health, Ithaca, NY, 14853

## Abstract

Yeast is commonly utilized in molecular and cell biology research, and *Yarrowia lipolytica* is favored by bio-engineers due to its ability to produce copious amounts of lipids, chemicals, and enzymes for industrial applications. *Y. lipolytica* is a dimorphic yeast that can proliferate in aerobic and hydrophobic environments conducive to industrial use. However, there is limited knowledge about the basic molecular biology of this yeast, including how the genome is duplicated and how gene silencing occurs. Genome sequences of *Y. lipolytica* strains have offered insights into this yeast species and have facilitated the development of new industrial applications. Although previous studies have reported the genome sequence of a few *Y. lipolytica* strains, it is of value to have more precise sequences and annotation, particularly for studies of the biology of this yeast. To further study and characterize the molecular biology of this microorganism, a high-quality reference genome assembly and annotation has been produced for two related *Y. lipolytica* strains of the opposite mating type, strain E122 (*MATA*) and 22301-5 (*MATB*). The combination of short-read and long-read sequencing of genome DNA and short-read and long-read sequencing of transcript cDNAs allowed the genome assembly and a comparison with a distantly related *Yarrowia* strain.

## INTRODUCTION

Y. lipolytica is an ascomycete yeast belonging to the class Saccharomycetes, known for its remarkable lipolytic and proteolytic capabilities, which enable it to thrive in hydrophobic, nutrient-rich environments [1]. Yarrowia therefore proliferates in environments rich in lipids and proteins, such as meat and dairy products, particularly fermented ones like cheeses and dry meats, as well as sewage or oil-polluted waters [2–4]. Yarrowia is significantly different from other hemiascomycetous yeasts in terms of its genomic features. For instance, it is a heterothallic yeast with two distinct mating types, MATA and MATB, and most natural isolates of this yeast are predominantly haploid [5]. Additionally, the G/C content in Y. lipolytica is notably high, averaging 49% and reaching nearly 53% in genes, particularly compared to the commonly used Saccharomyces cerevisiae with a genome containing 38.3% G/C [6]. Y. lipolytica and S. cerevisiae are estimated to have a common ancestor that existed 300 million years ago [7], making this pair of yeasts attractive for comparing the evolution of fundamental biological processes such as genome function and DNA replication. Yarrowia also has an unusually high number of intron-containing genes compared to S. cerevisiae and its related species [8]. The number of genes in Yarrowia is within the typical range for hemiascomycetous yeasts, but its genome size is 1.7 times larger than that of S. cerevisiae, which contains approximately the same number of genes.

Comparisons of the potential mechanisms of initiation of DNA replication in the budding yeasts, notably the predictions of DNA sequence-specific origins of DNA replication, have implied major differences between *Y. lipolytica* and *S. cerevisiae* [9]. *S. cerevisiae* and a small clade of highly related budding yeasts have origins of DNA replication that are composed of a number of DNA sequence-specific elements that constitute a functional origin of DNA replication [10,11]. The origin DNA elements in *S. cerevisiae* are recognized by the Origin Recognition Complex (ORC) and Cdc6, and two essential initiator subunits of ORC confer base-specific interactions via an inserted alpha-helix in the Orc4 subunit (Orc4-IH) and a loop in the Orc2 subunit (Orc2-loop) [9,12]. In contrast, *Yarrowia*, like nearly all other fungi and all animals and plants, lack these DNA sequence-specific recognition domains in Orc2 and Orc4, suggesting that it may have a more relaxed DNA sequence-specificity at its origins of DNA replication. Moreover, *S. cerevisiae* has lost RNA interference (RNAi) mechanisms but has gained Silent Information Regulator (SIR) proteins (Sir1, Sir3, and Sir4) that function in gene silencing and suppression of recombination of repetitive DNA sequences such as ribosome DNA (rDNA) and telomeric DNA [13–15]. Interestingly, *Y. lipolytica* lacks both RNAi and SIR-dependent gene-silencing proteins, including the RNAi silencing proteins such as Dicer and Argonaute (Ago) and SIR proteins, except for Sir2 [9]. All eukaryotes harbor the Sir2 gene that encodes an NAD-dependent histone deacetylase [16]. The SIR mechanism for suppressing recombination and silencing gene expression is particularly relevant in *S. cerevisiae* and its related budding yeasts such as *Kluyveromyces lactis,* where DNA-bound ORC interacts with SIR proteins and contributes to silencing the mating type genes and to the stability of rDNA and telomeric repeats [17–21]. It is, therefore, not known how *Y. lipolytica* silences gene expression or suppresses recombination of repetitive rDNA and telomeric DNA.

One possible explanation for the occurrence DNA sequence-specific origins of DNA replication in some budding yeast species, such as *S. cerevisiae*, is that these organisms have lost much of the intergenic DNA and lack introns, therefore they possess a very gene-dense genome relative to their genome size. The presence of DNA sequence-defined origins in gene-rich organisms, such as *S. cerevisiae,* could provide an advantage in recruiting ORC to intergenic sites within these species, thereby avoiding conflicts between DNA transcription and the initiation of DNA replication, which can result in genome instability [9,22]. As a result, organisms like *S. cerevisiae*, with high gene density and smaller intergenic regions, have evolved an efficient mechanism to ensure that the replication complex can find appropriate sites for initiation and avoid initiating DNA replication in a transcribed region. By gaining a deeper understanding of how DNA replication is initiated in a variety of species, including human cells and in diverse yeasts such as *Y. lipolytica*, further insights into how origins of DNA replication are located in the genome and the replication strategies in eukaryotic cells will become apparent [23]. For this reason the complete genome sequences and assemblies of two *Y. lipolytica* strains of opposite mating type were performed to assist in subsequent studies of whole genome DNA replication and gene silencing mechanisms. Both long-read sequencing using PacBio and Oxford nanopore methods and short-read Illumina-based methods of both genomic DNA and cDNA were used to generate and assemble the genome of the two *Y. lipolytica* strains.

Previously, the main reference genome for *Y. lipolytica* was strain CLIB122 (also called E150), a derivative from a mating between a French isolate W29 and an American isolate YB423-12 (CBS 6124-2) (see Figure 1), that was obtained through short read Sanger sequencing [6,24]. A number of other strains have been shot-gun sequenced and their genomes compared, resulting in a recent summary of these genome comparisons [25]. The estimated size of the six-chromosome genome was ∼21 Mb [24]. However, these assemblies contained gaps, and the telomeric ends as well as rDNA repeats could not be integrated into the genome assembly due to their repetitive nature. The CLIB122/E150 strain and its derivatives, including the commonly used PO1f strain (Figure 1) [26], are the main strains used for laboratory studies. A summary of the parental backgrounds, alternative strain names and genotypes is shown in Table 1, including the *MATA* and *MATB* strains employed herein. A high-quality, near-contiguous genome assembly of a distantly related *Y. lipolytica* strain DSM 3286 (a German strain) was obtained using a combination of long-read and short-read genomic DNA sequencing [27]. This allowed the characterization of the repetitive rDNA and telomeric regions and the observation that rDNA clusters are located near the telomeric regions. Additionally, the genetic and phenotypic diversity of 56 haploid strains of *Y. lipolytica* was investigated by sequencing of a diverse set of *Y. lipolytica* strains collected from various geographical and biological origins, and included revision of the version of the E150 *Y. lipolytica* strain genome sequence and annotation [25].

**Figure 1:**
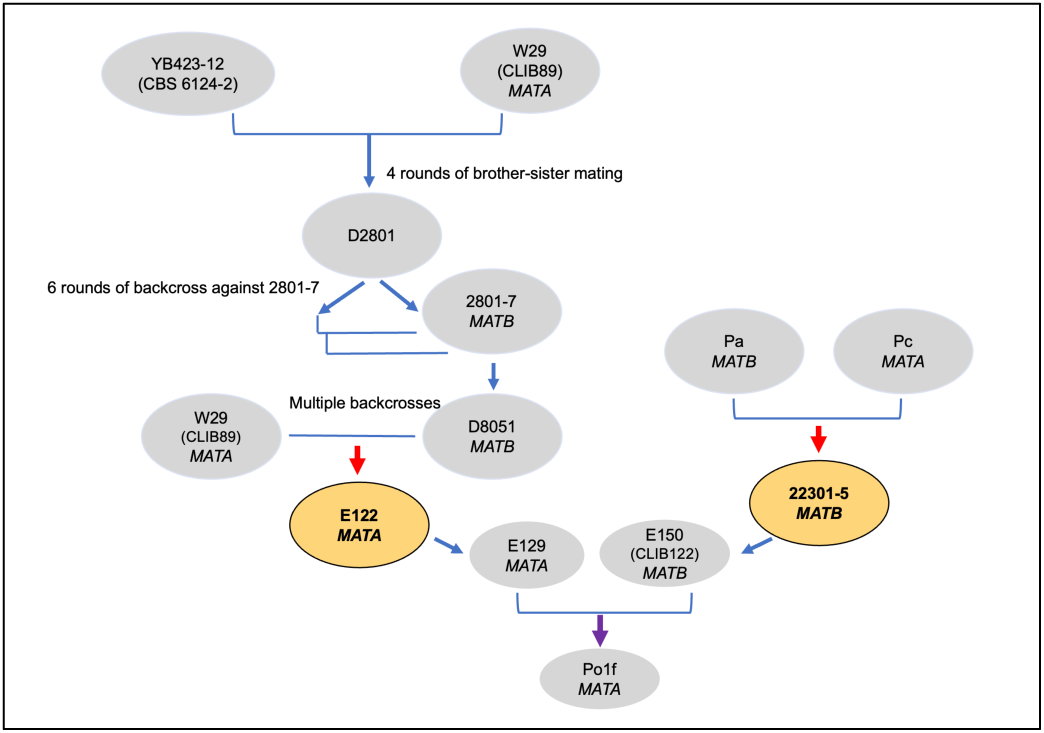
Breeding and backcrossing strategy for strain development in *Yarrowia lipolytica*. The flowchart illustrates the lineage and mating strategy employed to derive key strains of *Y. lipolytica*. YB423-12 lys1.13 and W29 (CLIB89 *MATA*) underwent four rounds of brother-sister mating to generate the intermediate strain D2801. D2801 was subjected to six rounds of backcrossing against 2801-7 *MATB*, resulting in the development of the strain D8051 *MATB*. Parallel strategies were employed using Pa *MATB* and Pc *MATA*, which were crossed to form 22301-5 *MATB*. E122 *MATA* was derived from multiple backcrosses and further developed into strains like E129 *MATA*, E150 (CLIB122 *MATB*), and Po1f *MATA*, widely used for research and industrial applications. Color-coded ovals indicate key final strains derived from these processes (e.g., E122 *MATA* and 22301-5 *MATB*). Blue arrows represent mating and backcrossing steps; red arrows indicate the lack of a *Ura3* marker, and purple arrows indicate the lack of *Xpr2* and *Axp* genes.

**Table 1.**
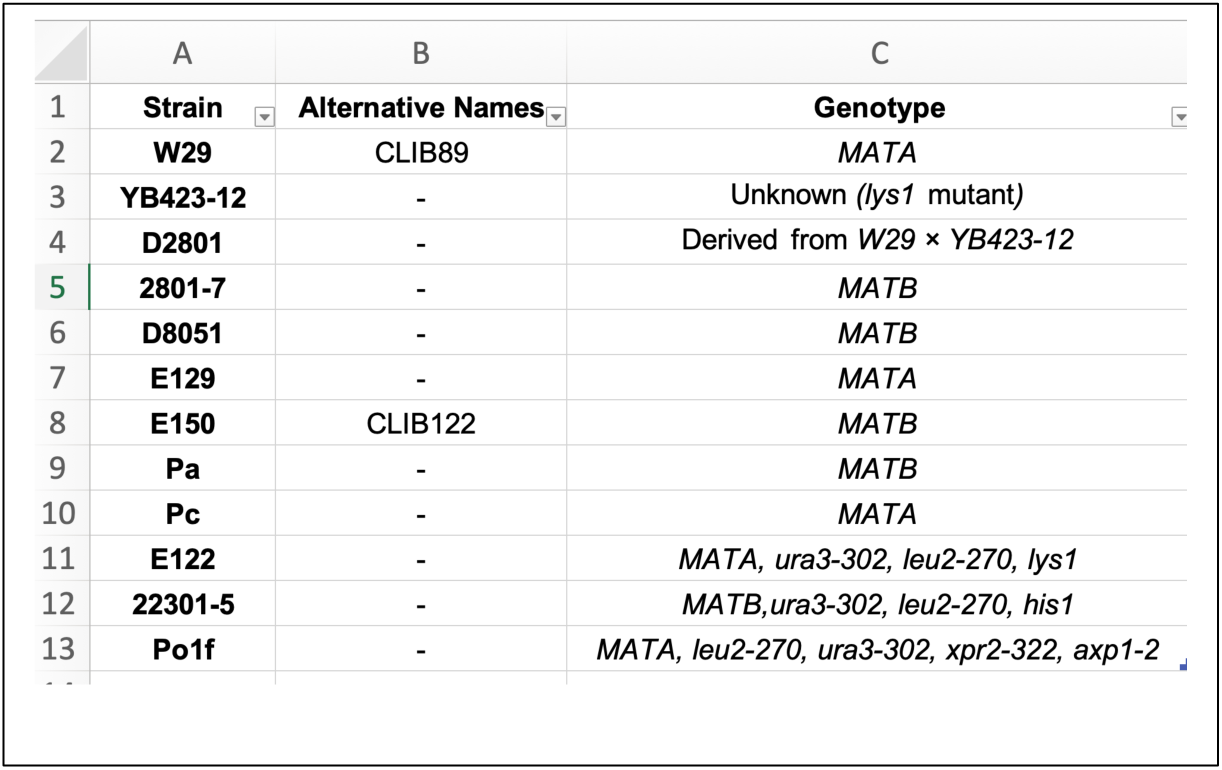
Summary of *Yarrowia lipolytica* Strains from Figure 1. This table provides an overview of all *Yarrowia lipolytica* strains referenced in Figure 1, including their alternative names and genotypes. While only the E122 *MATA* and 22301-5 *MATB* strains were used for experimental work in this study, the table summarizes the parental origins from which these strains were derived, along with their respective genotypes. The genotype of E122 has been changed from the literature [57] to *MATA*, *ura3-302, leu2-270, lys1* (see results).

In this study, we have used both long and short-read sequencing of genomic DNA and cDNA copies of RNA transcripts to precisely compare the genomes of genetically related *MATA* and *MATB* strains that are distant from the DSM 3286 strain isolated in Germany and other geographically diverse strains that have been sequenced [25,28]. Chromosomal rearrangements were observed comparing the multiple isolates. The sequences allowed annotation of the genome and revealed the presence of many repeated DNA sequences such as transposable elements, including LTR-retrotransposons, LINE elements, and DNA transposons from various families, which are distributed variably among strains. Moreover, the distribution of and number of rDNA repeats was analyzed.

## MATERIAL AND METHODS

Strains: Two related *Yarrowia lipolytica* strains of opposite mating type were obtained from Richard A. Rachubunski, University of Alberta, Canada, and single clones were isolated and used for both genome and transcript sequencing. The strains were 22301-5 (*MATB*) and E122 (*MATA*, alternatively called CLIB120) [5,29,30] (Figure 1 and Table 1). E122 is *MATA*, *ura3-302*, *leu2-270*, *lys8-11(*but see results for *lys-8)* and is related to the *MATB* strain E150 (CLIB122 and is the current DNA sequence reference strain) [6,24]. Strain 22301-5 is from a different lineage and is *MATB*, *his-1, ura-302, leu2-270,* (Figure 1 and Table 1).

### DNA and RNA Preparation and Sequencing

#### RNA

To isolate the high molecular weight RNA, the TRIzol Plus RNA Purification Kit from Thermo Fisher Scientifics was used. Briefly, 5ml liquid of yeast culture was grown to an OD^600^ of 2.0 and cells were pelleted and lysed with TRIzol™ reagent according to the user manual. Following lysis, the RNA present in the sample was bound to the PureLink RNA Mini Kit Spin Cartridge (12183018A, Invitrogen) where it was washed to remove contaminants. Lastly, eluted RNA was stored in 50-microliter aliquots at −80°C. To generate a short-read RNA, a Direct-zol RNA purification kit from Zymo Research (Cat # R2050) was used according to the user manual. Long-read sequencing of RNA transcripts was performed as follows. The RNA was prepared using the ONT SQK-PCS109 kit according to the manufacturer’s instructions, and it was loaded onto a PromethION P24 system with a PROM-0002 flow cell. Base-calling was performed using the live hac base-calling guppy version 3.2.10. Two cells of *MATA* and *MATB* were run for each experiment.

#### DNA

For the generation of high molecular weight DNA and ultra-long nanopore sequencing reads, cells in a 100 ml culture grown overnight at 30°C in Yeast extract, Peptone and Dextrose (YPD) were harvested, washed in sterile distilled water and incubated for 2 hr at 37°C in 10 ml SEB buffer (0.9 M sorbitol, 0.1 M EDTA, 0.8% β -mercaptoethanol) containing 5 mg Zymolyase 20T (Sunrise Science products, CAT#N0766391). Protoplast formation was monitored by phase contrast microscopy. The protoplasts were then harvested, resuspended in 3 ml TE Buffer (Tris-EDTA, pH 8.0), then 300 μl 10% SDS was added, and the samples were incubated at 65°C for 30 minutes. 1 ml of 5 M potassium acetate was added, and the samples were kept on ice for 1 h. The supernatant was recovered after centrifugation, and DNA was precipitated by adding 0.1 volume of 3 M sodium acetate and 2.5 volumes of ethanol at −20°C for at least 1 h. The DNA was recovered by centrifugation and resuspended in 3 ml TE [31]. Then 100 µg/mL of proteinase K along with 50 µg/mL RNase A were added and the samples were incubated at 37°C for 3 hrs. After centrifugation for 45 min at 12,000 × g and 4°C, the supernatant was collected and transferred to a 2-ml Eppendorf tube. Samples were then extracted two more times with phenol/chloroform/isoamyl alcohol and one final time with chloroform. To precipitate DNA, 2-2.5 volumes of 100 % ice-cold ethanol was added to the aqueous phase along with 1/10 volume of 3 M sodium acetate, mixing by inversion, and samples were incubated at −20°C for at least 1 h. The DNA was recovered by centrifugation for 20 min at 12,000 × g and 4°C, and the pellet was subsequently washed three times with 2 ml of 80% (vol/vol) ethanol. The pellet was then air dried and dissolved in 100 μl of Tris-EDTA.

DNA fragment length and molecular weight distributions of genomic DNA samples were evaluated using a Femto Pulse pulse-field capillary electrophoresis system (Agilent). More than 5ug of DNA was size selected via SRE XS (Circulomics). The full reaction was repaired and end prepped with NEBNext FFPE DNA Repair Buffer and Ultra II End prep kit (NEB). The reaction was cleaned up with 1X Ampure beads and precipitated with ethanol. DNA was bound to ONT adapter from the SQK-LSK109 kit (ONT) via NEBnext Quick T4 ligation module (NEB). DNA was resuspended in SQB buffer (ONT) and loading beads (ONT) and sequenced on one PromethION 24 cell PROM0002 with a three-day run time.

For short-read DNA sequencing, the YeaStar Genomic DNA Kit from Zymo Research (CAT# D2002) was used according to the user manual. DNA sequencing libraries were prepared per the manufacturer’s instructions with a Kapa DNA hyperprep kit (Roche CAT #KK8504). It was loaded on an Illumina MiSeq with a PE150 v2 format.

### DNA Sequence Assembly

#### Long-Read processing

The unprocessed long-reads were produced using Guppy v.5 base-caller from Oxford Nanopore Technologies (https://github.com/nanoporetech). To assemble the reads, the long-read assembly pipeline Flye v. 2.9-b1774 (https://github.com/fenderglass/Flye) [32] was used in nano-hq mode, which is intended for high-quality reads (<5% error rate). The minimum overlap between reads was set to 7KB. The pipeline was run with five iterations of polishing.

#### Short-Read processing

The paired-end reads were trimmed using Cutadapt v.3.7 [33]. Cutadapt removes adapter sequences from high-throughput sequencing reads [33]. BWA v.0.7.17-r1188 [34] was used to index the long-read assembly and align the trimmed short-reads to the assembly. The alignments were sorted and indexed using Samtools v.1.14 [35]. Pilon v1.24 (https://github.com/broadinstitute/pilon) [36] was used for polishing the long-read assembly with the aligned short-reads. We obtained exactly one contig per chromosome and mitochondria for *MATB* and an extra contig for *MATA*. To scaffold the extra contig, we used RaGOO [37] and the assembled *MATB* as the reference genome sequence.

### Transcript assembly

For the transcriptome assembly, three different transcriptome assemblies were combined: one from the short-read sequencing, a second one from the Nanopore long cDNA reads, and a third combining both. The short-reads were first trimmed using Trimmomatic v.0.38 [38]. The trimmed reads were aligned to the genome using hisat2 (v.2.2.1). A short-read only transcriptome was then assembled using Stringtie v.1.3.6 [39]. The long cDNA reads were *de novo* assembled using Oxford Nanopore’s Workflow Transcriptomes (wf-transcriptomes) pipeline (v1.1.1). A third transcriptome that used long- and short-reads was assembled using TASSEL [40] (https://github.com/kainth-amoldeep/TASSEL). These three transcriptomes were then combined using gff compare (v.0.12.2) [41] and the output was used as input to the annotation pipeline.

### Gene annotations

The MAKER-P (v.3.0) [42] pipeline was used to annotate protein-coding genes in the two strains 22301-5 (*MATB*) and E122 (*MATA*). As evidence, we used all annotated proteins from *Yarrowia lipolytica* (budding yeasts) downloaded from the NCBI protein database. These protein sequences were clustered using CDHit-est (v4.6) [43] with parameters (-c 0.95 -n 10 -d 0 -M 3000 -t 1). For transcript evidence, the combined transcriptome from gff compare was used for transcript assembly. The assembled transcripts were checked and filtered for intron retention using Suppa (v2) [44]. For gene prediction, we used Augustus (v3.3) [45,46] trained on *Y. lipolytica* and FGENESH (http://www.softberry.com) trained on *S. pombe*, respectively. Repeat masking was initially performed using Repeatmasker (RepeatMasker Open-4.0) with the Ensembl repeat annotation pipeline, employing parameters (-nolow -gccalc -species “Fungi” -engine ncbi) and a yeast specific library. The resulting repeat-masked genomes were then used for annotation with MAKER-P, incorporating additional protein evidence. To improve repeat identification and capture lineage-specific elements not represented in curated libraries, we additionally performed *de novo* repeat annotation using the EDTA tool [47]. This combined approach allows for more comprehensive detection and classification of repetitive elements. Additional improvements to structural annotations were done using PASA (v2.3.3) [48] using the assembled transcriptome and fungal EST from NCBI using query (EST[Keyword] AND fungi [Organism]. Gene identifiers were assigned using existing nomenclature schema established for *Yarrowia* for each strain [6,24]. Functional domain identification was completed with InterProScan (v5.38-76.0) [49]. TRaCE [50]___was used to assign canonical transcripts based on domain coverage, protein length, and similarity to transcripts assembled by StringTie (v1.3.4a) [39]. Finally, the gene annotations were imported to Ensembl core databases, verified, and validated for translation using the Ensembl API [51].

Genome comparative analysis was done using 5 *Yarrowia lipolytica* strains including 15 closely related species and outgroups providing the foundation for building protein-based gene trees based on the EnsemblCompara pipeline [52].

## RESULTS

A hybrid assembly approach was employed by pairing Illumina short-read DNA sequences and high-quality long-read DNA sequences with much-improved base-calling using NANOPORE Technologies’ Guppy 5 base caller with very high mean coverage, as indicated in Figure 2A and Table 2. Longer contiguous and quasi-contiguous genome assemblies for the E122 *MATA* and 22301-5 *MATB* genomes were obtained, compared to previous assemblies of the related French isolates. The improvements over the reference assembly for strain CLIB122-E150 included the incorporation of telomeric repeats on each chromosome, as well as a decrease in the estimated number of missing essential gene markers according to BUSCO assessment. We assessed the quality of the assemblies using BUSCO version 4.1.1 [53], which employs AUGUSTUS as the gene predictor in genome mode on the *Saccharomycetes* lineage set of 2137 essential genes. The results indicate a completeness index of 96.8% (Figure 2B). *Yarrowia* appears to be missing genes that are present in other *Saccharomycetes,* or the genes are diverged enough so that BUSCO does not detect them.

**Figure 2.**
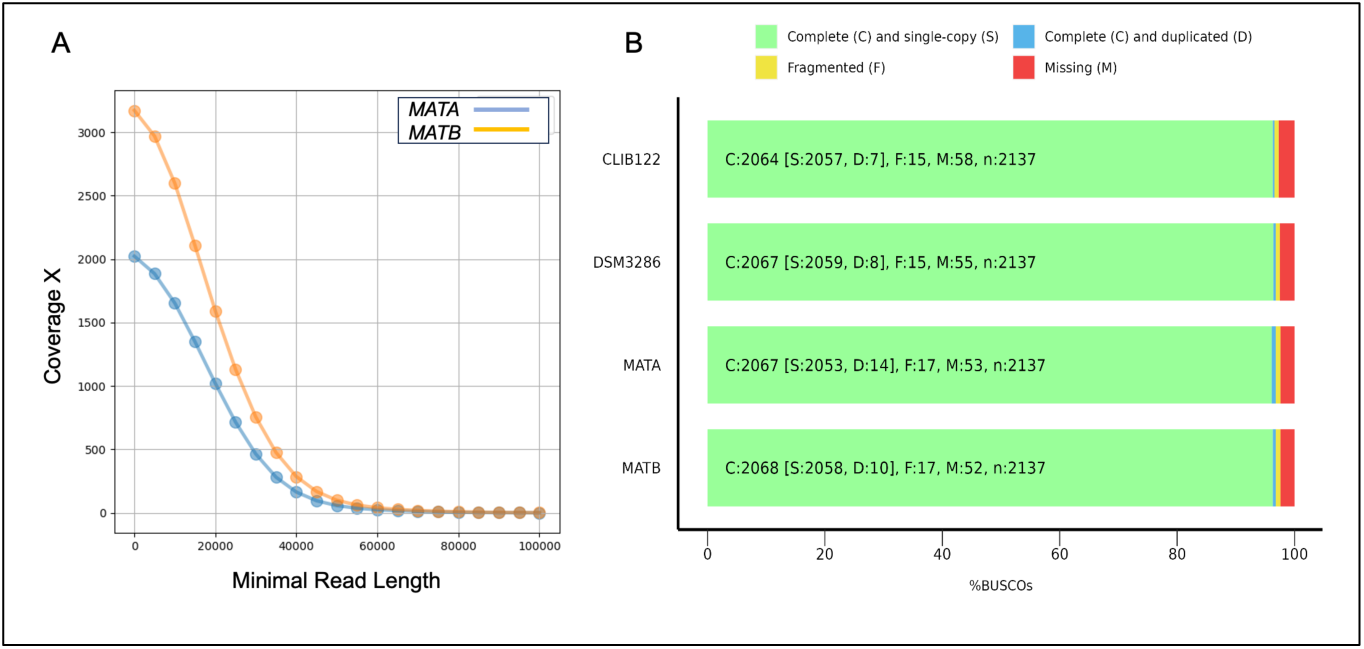
Comparison of genome assembly metrics between *Yarrowia lipolytica* strains E122 (*MATA*) and 22301-5 (*MATB*) **A.** Coverage versus minimal read length distribution: Coverage (X) is plotted against minimal read lengths for both strains. The blue line represents E122 *MATA*, while the orange line represents 22301-5 *MATB*. The higher coverage for 22301-5 *MATB* indicates a more profound sequencing effort compared to E122 *MATA*. The gradual decline in coverage with increasing read length reflects the expected distribution of read sizes. **B.** Figure produced by BUSCO depicting genome completeness analysis across four *Yarrowia lipolytica* strains (CLIB122, DSM 3286, E122 *MATA*, and 22301-5 *MATB*). C: Number of complete BUSCO genes, divided into S: Single-copy genes and D: Duplicated genes. F: Number of fragmented BUSCO genes. M: Number of missing BUSCO genes. N: Total number of BUSCO groups analyzed (2137). Each bar represents the percentage distribution of these categories for a strain. Most BUSCO groups are complete and single-copy (light green), reflecting high genome assembly quality. Slight variations in duplicated (blue), fragmented (yellow), and missing (red) categories highlight subtle differences in genome assemblies among the strains.

**Table 2:** Comparative genome assembly statistics for *Yarrowia lipolytica* strains E122 (*MATA*) and 22301-5 (*MATB*) Total Read Length: Total bases sequenced for each strain; Mean Coverage: Average sequencing coverage (sequencing depth); Reads N50/N90: Median (N50) and 90th percentile (N90) read lengths; Total Length: Total assembled genome length in base pairs; Fragments N50: Fragments: Number of assembled fragments; Largest Fragment: Largest assembled fragment in base pairs; Total Length (Alternate): Total length in base pairs across all fragments. These statistics provide a detailed comparison of genome assembly quality, highlighting the structural integrity of the two *Yarrowia lipolytica* strains.

|  | **E122 MATA** | **22301-5 MATB** |
| --- | --- | --- |
| Total Read Length | 35055999261 | 53578162586 |
| Mean coverage | 1630 | 2364 |
| Reads N50/N90 | 19835 / 6596 | 19758 / 6756 |
| Total Length | 21019611 | 21008502 |
| Fragments N50 | 3712330 | 3688210 |
| Fragments | 8 | 7 |
| Largest fragment | 4317224 | 4320808 |
| Total length | 20313536 | 21008502 |

A dot plot produced with chromeister (release 1.5.a.) [54] compared the E122 *MATA* and 22301-5 *MATB* genomes, revealing very high similarity between these two related genomes, as expected (Figure 3A). The similarity score indicated a divergence of just 0.003. In contrast, a similar comparison between 22301-5 *MATB* and the German isolate DSM 3286 showed considerable genome rearrangements and a reduced similarity score of 0.0031 (Figure 3B), as noted previously [27]. When the genes that were expressed in E122 *MATA* but not 22301-5 *MATB* were analyzed, they were in and surrounding the *MATA* locus and included *Sla2,* encoding an actin binding protein, and *Apn2,* encoding a DNA-(apurinic or apyrimidinic site) lyase, both genes that flank the *MATA* locus [55,56], as well as the *MATA1* and *MATA2* mating type genes [56].

**Figure 3.**
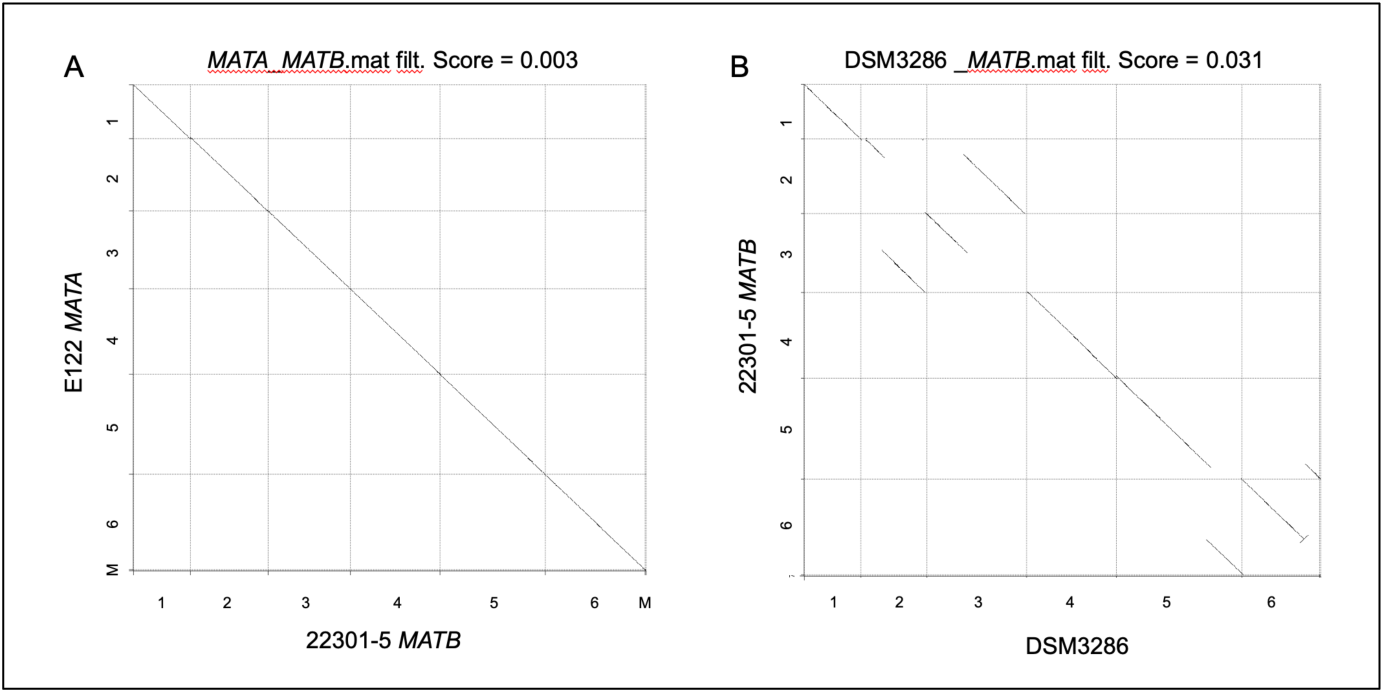
Assembly alignment comparison. **A.** Dot plot of the final assemblies for E122 *MATA* and 22301-5 *MATB* shows synteny between the two genomes. The diagonal alignment highlights the high degree of sequence conservation, with a calculated score of 0.003, indicating minor structural or sequence variations between the two genomes. This data highlights the quality of sequencing and genome assembly while comparing structural differences between these two strains of *Y. lipolytica*. **B.** Dot plot of the sequences for 22301-5 *MATB* and DSM 2386. The diagonal alignment highlights the genome rearrangements between the two strains, thereby reflecting a lower calculated score of 0.031.

The genotypes of the two strains cited in the literature are E122, *MATA*, *ura3-302, leu2-270, lys8-11* and 22301-5 *MATB, ura3-302, leu2-270 and his1.* We confirmed the auxotrophic markers on selective media and analyzed the DNA sequences of the *URA3*, *LEU2*, *HIS1* and *LYS* genes. The *URA3, HIS1* and *LEU2* gene sequences showed mutations in the relevant strains, however we found that E122 had a deletion in the *LYS1* gene encoding homocitrate synthase that could explain the lysine autotrophy. The *LYS1* gene was wild type in strain 22301-5. The first report of the lys8-11 mutation in E122 we could find was reported by Nuttley et al. [57], however a description of the mutation was not provided. We therefore suggest that strain E122 is *lys1* and not *lys8* and we hence forth will refer its genotype as *MATA*, *ura3-302, leu2-270, lys1* (Table 1).

### Gene annotations and comparative analysis

The structural gene annotation pipeline identified 7,728 and 7,769 genes in *Yarrowia lipolytica* strains E122 (*MATA*) and 22301-5 (*MATB*), respectively (Table 3). This gene count surpasses that of previously reported *Yarrowia* strains DSM 3286 [27] with 6,439 protein-coding genes and the reference strain CLIB-122 [58] with 6,448 protein-coding genes. To further evaluate annotation quality, we utilized the Annotation Edit Distance (AED) score generated by MAKER-P [42]. An AED score of 0 indicates genes supported by evidence, while a score of 1 indicates a lack of evidence. Employing mRNA and homology evidence, as described in the methods, to calculate AED scores yielded 6,297 and 6,212 genes with some evidence (AED score < 1) in strains E122 *MATA* and 22301-5 *MATB*, respectively. In contrast, strains DSM 3286 and CLIB-122 (E150) exhibited 6,236 and 6,387 protein-coding genes with AED scores <1 (Table 3).

**Table 3:** Comparative Gene and Transcriptomic Features of *Yarrowia lipolytica* Strains. This table presents the genomic and transcriptomic characteristics of four *Yarrowia lipolytica* strains (E122 *MATA*, 22301-5 *MATB*, DSM3286, and CLIB122), highlighting differences in gene structure and composition. The *Yarrowia lipolytica* strains exhibit notable variations in gene structure, with 22301-5 *MATB* having the highest gene count (7,769) and DSM3286 the lowest (6,439), while E122 *MATA* has the longest median gene length (1,473 bp). The high proportion of single-exon genes (78.2–85.2%) and low exons per transcript (∼1.2–1.3) suggest a predominantly condensed genetic structure, with CLIB122 having the most compact gene architecture and E122 *MATA* showing greater structural complexity. This data provides insights into structural genomic variation and transcriptional complexity across these *Y. lipolytica* strains.

| Column1 | E122 *MATA* | 22301-5 *MATB* | DSM3286 | CLIB122 |
| --- | --- | --- | --- | --- |
| Total protein-coding genes | 7728 | 7769 | 6439 | 6448 |
| Gene count AED <1 | 6297 | 6212 | 6236 | 6387 |
| Gene length(median) | 1473 | 1434 | 1287 | 1245 |
| Exon count | 7975 | 7861 | 7661 | 7457 |
| Exon length(median) | 1071 | 1050 | 1033 | 1059 |
| Intron count | 1678 | 1649 | 1425 | 1070 |
| Intron length(median) | 208 | 203 | 193 | 207 |
| Peptide count | 6312 | 6221 | 6236 | 6387 |
| Peptide length(median) | 400 | 405 | 408 | 403 |
| Exons per transcript(Avg) | 1.3 | 1.3 | 1.2 | 1.2 |
| Single-exon gene count(%) | 78.2 | 78.3 | 81 | 85.2 |

When comparing annotation features between the strains E122 *MATA* and 22301-5 *MATB* using genes filtered by AED <1, it was observed that most protein-coding genes contained a single exon. Moreover, we found more genes in E122 *MATA* and 22301-5 *MATB* with multiple introns compared to the previously analyzed strains (Figure 4A). The average intron size decreased with introns closer to the 3’-end of the gene (Figure 4B). The shortest introns were ∼40 base pairs and the longest intron was 6782 base pairs (Figure 4C and D). The median gene lengths in E122 *MATA* and 22301-5 *MATB* were higher than previously estimated, reflecting their increased intron counts compared to the other two strains (Table 3). *Yarrowia lipolytica* genomes are known to be intron-rich, with previous estimates of 15% of genes containing introns, which is 4 times that of *S. cerevisiae* [8]. Intron-containing genes in E122 *MATA*, 22301-5 *MATB* and the DSM 3286 strain represented ∼20% of the protein-coding genes, highlighting the improvement in genome assembly using long-read technology compared to CLIB-122. 80% of these intron-containing genes were mono-intronic, compared to 20% that were multi-intronic (with up to five introns). The internal exons of the multi-intronic genes were mostly short compared to 1st intron (Figure 4B and D).

**Figure 4.**
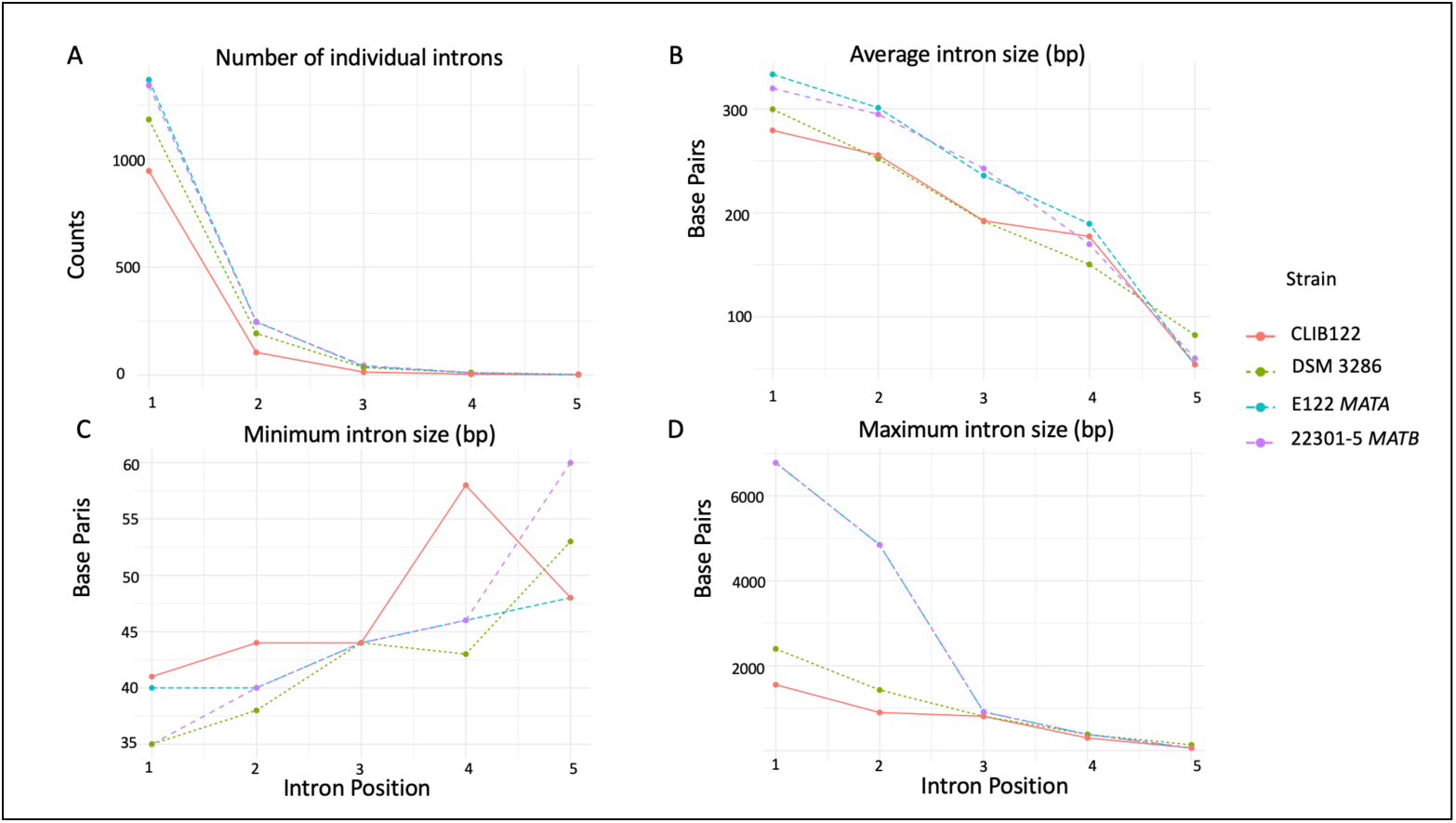
Comparative analysis of intron properties across *Yarrowia lipolytica* strains CLIB122, DSM 3286, E122 *MATA*, and 22301-*5 MATB* strains. **A.** Number of Individual Introns is shown for each intron position in all genes. **B.** The average intron size in base pairs for each intron position in all genes. **C.** The minimum size of introns in base pairs at each intron position. **D.** The maximum intron size in base pairs for each intron position in all genes.

A total of 9,453 orthologous genes were identified across the four *Yarrowia lipolytica* strains analyzed (E122 *MATA*, 22301-5 *MATB*, DSM 3286, and CLIB-122; Figure 5A). Of these, 6,050 were core genes found in all four strains, as indicated by the largest bar (pattern “1111”) in the UpSet plot (Figure 5B, lower panel). This number closely matches the 6,042 core genes previously detected in a survey of seven *Y. lipolytica* strains and is slightly lower than the 6,528 core genes found among 54 strains in a pan-genome analysis (25). This modest difference between the core and pan-genome further supports the low genetic diversity of *Y. lipolytica*, consistent with prior reports (25). We also identified 1,204 gene ortholog groups that are uniquely present in both laboratory strains (E122 *MATA* and 22301-5 *MATB*) but absent from DSM 3286 and CLIB-122, as shown by the bar corresponding to the “1100” intersection pattern in Figure 5B. To further understand the potential functions of these 1,204 unique ortholog groups, we performed Gene Ontology (GO) enrichment analysis. Figure 5C displays the significantly enriched molecular functions, and Figure 5D shows the enriched biological processes. Gene Ontology (GO) analysis of the 1,204 unique ortholog groups revealed only 14 genes with associated GO terms. Among these, enrichment analysis indicated significant overrepresentation of functions related to iron and sulfur cluster binding (GO:0051537), methylmalonate-semialdehyde dehydrogenase (acylating) activity (GO:0004491), oxidoreductase activity (GO:0016491), and TBP-class protein binding (GO:0017025) (Figure 5C). Biological process enrichment further highlighted DNA-templated transcription initiation (GO:0006352) (Figure 5D).

**Figure 5.**
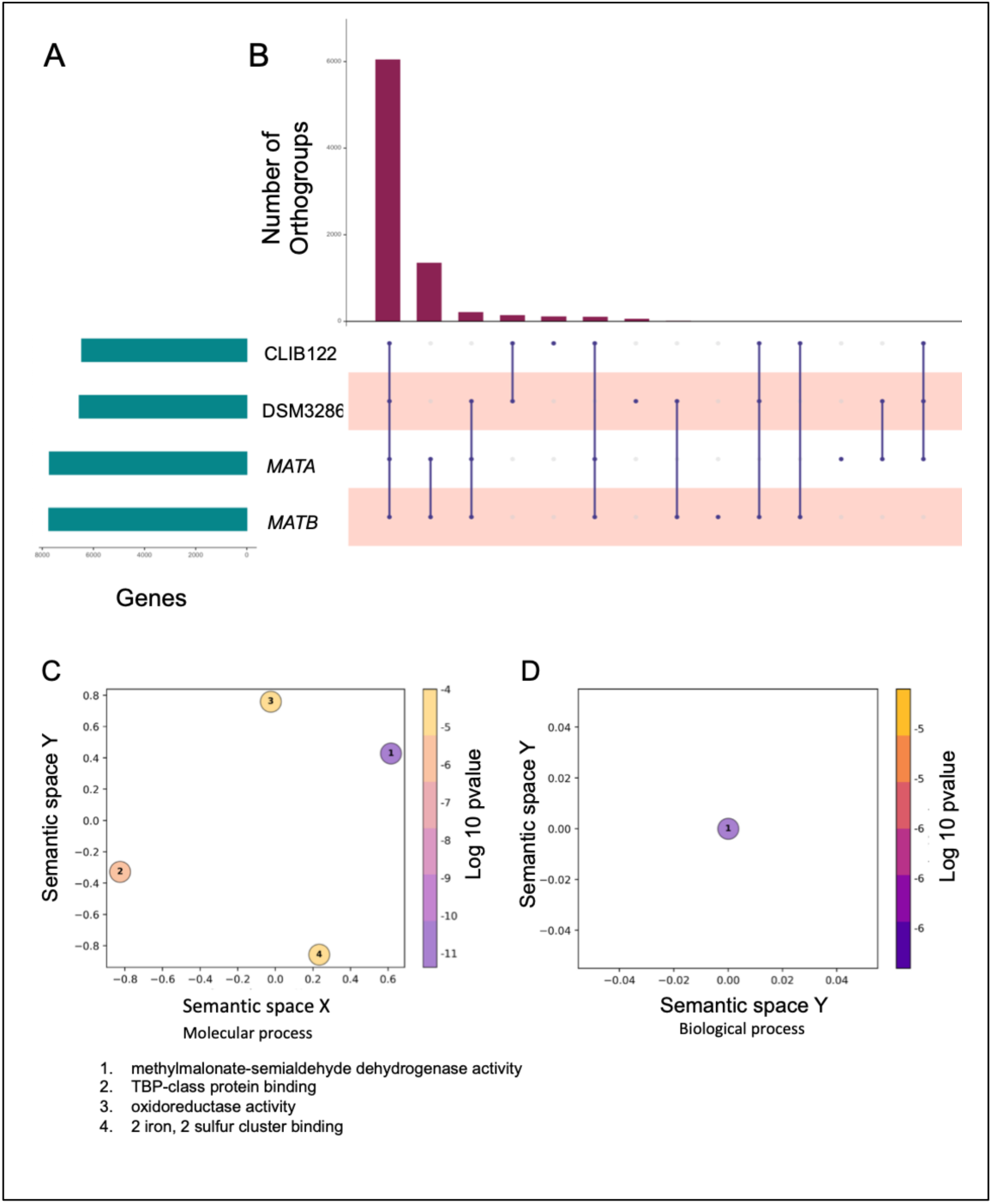
Ortholog distribution and functional enrichment in four *Y. lipolytica* strains. **A.** Total predicted genes per genome (turquoise bars). **B.** UpSet plot of ortholog-group intersections. Dot matrix shows presence (blue) or absence (grey) of each strain; maroon bars give the size of each intersection. The tallest bar (pattern 1111) represents 6,050 core orthologs shared by all strains; the next bar (1100) marks 1,204 orthologs found only in E122 *MATA* and 22301-5 *MATB*. **C.** Molecular-function GO terms enriched in the 1,204 orthologs. Circle color denotes statistical significance (-log10 *p-*value) end lists the enriched functions. **D.** Biological-process GO enrichment for the same set; only DNA-templated transcription initiation is significant.

### Analysis of Repeat Sequences

We compared repeat landscapes in four *Y. lipolytica* genomes using a yeast-specific RepeatMasker library.(Figure 6A). The E122 (*MATA*) and 22301-5 (*MATB*) assemblies were built entirely from high-quality ONT long reads polished with Illumina data (Flye v2.9; five polishing rounds). In contrast, the DSM3286 reference derives from a hybrid ONT + Illumina workflow (Canu v1.6 → Nanopolish → 19× Pilon) [27], while the legacy CLIB122 genome was produced by large-insert Sanger shotgun sequencing [8]. E122 and 22301-5 contain markedly more RNA repeats, LTRs, LINEs, and SINEs than DSM3286, whereas CLIB122 shows the lowest counts in every retro-element class (Figure 6). Because short-read and Sanger assemblies routinely collapse multi-copy retro-transposons and leave telomere/rDNA gaps, we attribute most of this disparity to assembly bias rather than true biological loss of RNA elements. Within the long-read genomes, repeat profiles are concordant: RNA repeats ∼ 30 %, simple repeats 35-40 %, and LTRs 15-18 %. DSM3286 shows a modest skew toward simple repeats (∼55 %) and away from RNA repeats (∼10 %), consistent with partial collapse during hybrid polishing, a signature of short-read polishing workflows (e.g., multiple *Pilon* rounds) that collapse multi-copy retro-elements and inflate adjacent microsatellites [59]. Controlled benchmarks show that short-read assemblies miss 26–57 % of transposable-element (TE) insertions that long-read assemblies recover [60]. To complement the RepeatMasker-based annotation, we also performed *de novo* repeat identification using the EDTA tool [47] which captures previously unannotated and lineage-specific repeat elements not represented in curated libraries. A summary of the EDTA-identified repeats, including classification and abundance in both *MATA* and *MATB* strains, is provided in Figure 6B. To assess their functional impact, we analyzed overlaps between annotated repeats (from both curated and *de novo* sources and gene loci). This analysis revealed that 661 genes in the *MATA* strain and 649 genes in the *MATB* strain overlap with repeat elements, suggesting that repeats may influence gene regulation or genome architecture in *Y. lipolytica*.

**Figure 6.**
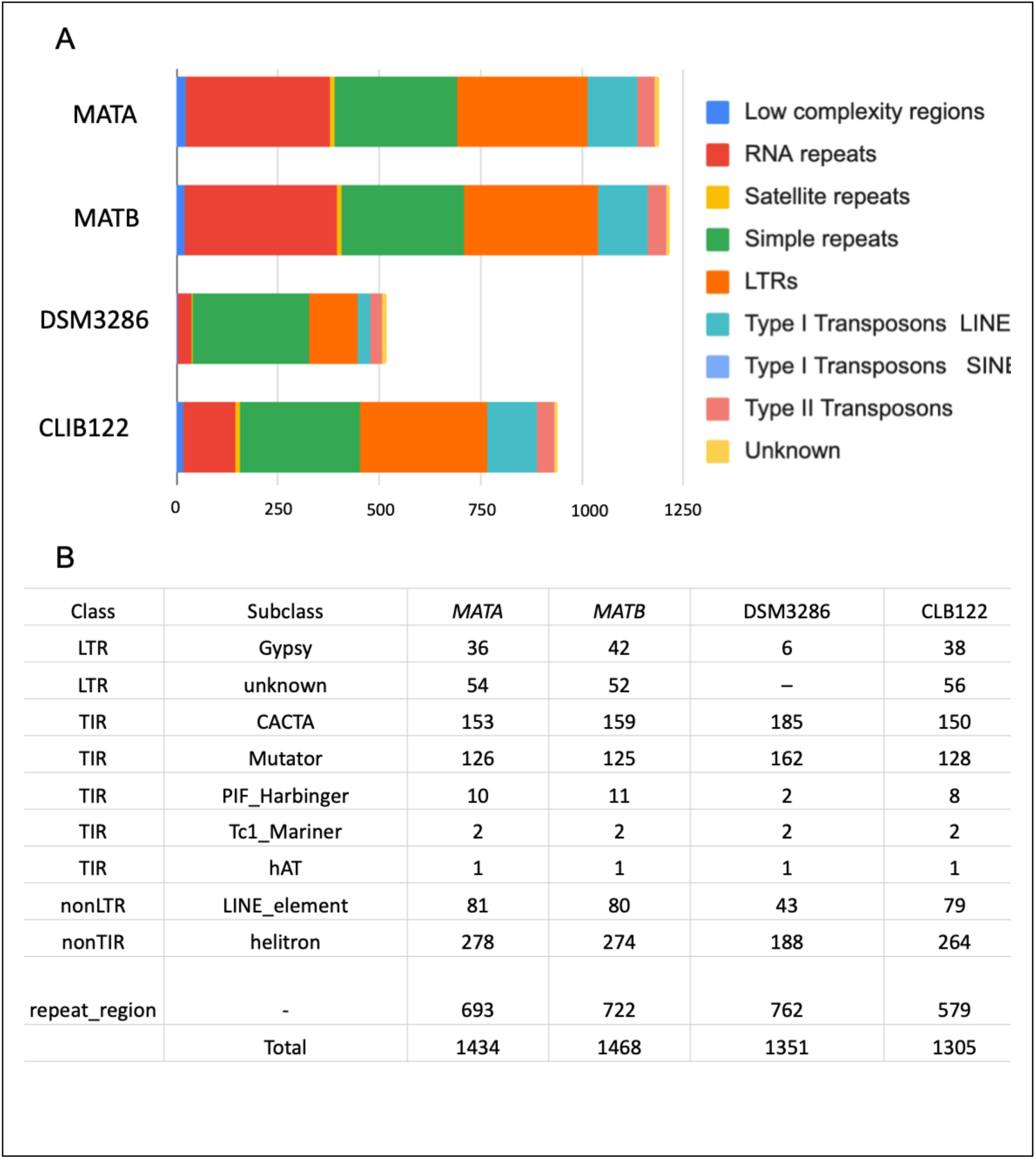
Comparative repeat composition in the genomes of *Yarrowia lipolytica* strains E122 *MATA*, 22301-5 *MATB*, DSM 3286, and CLIB122. **A.** Repeat element composition in four *Y. lipolytica* genomes. Stacked bars show absolute counts of major repeat types across E122 *MATA*, 22301-5 *MATB*, DSM 3286, and CLIB122. **B.** Classification of repeat elements identified by EDTA. Table summarizes the number of elements per transposon subclass in each genome, including LTRs, TIRs, non-LTRs, and helitrons.

As previously observed [27], the rDNA repeats consisting of the 18S and 28S genes were located at the ends of chromosomes B (right end), C (both ends), E (right end) and F (both ends), and lie adjacent to the telomeres (Figure 7A). The 5S rDNA genes are scattered throughout the genome on every chromosome. For the E122 *MATA* and 22301-5 *MATB* strains, the calculated size of these repeats in kilobase pairs (yellow bar) and the number of rDNA repeats (blue bar) for each region of the genome is shown in Figure 7B.

**Figure 7.**
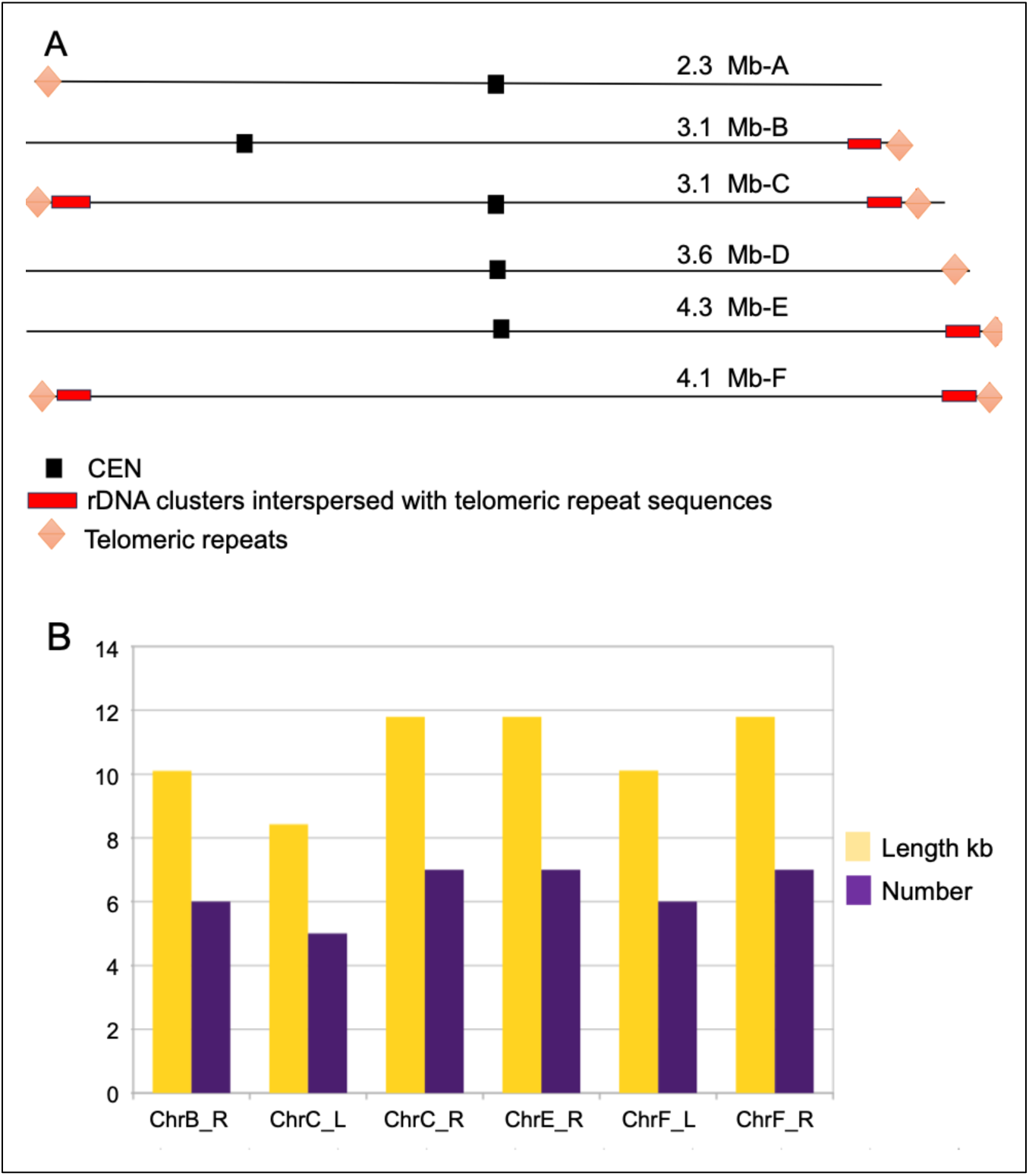
Analysis of rDNA Repeats. **A.** Genome landscape of six chromosomes of *Y. lipolytica* mapped rDNA (red bar) and telomere (orange diamond) sequences. The known centromere sequences are shown with black bars. **B.** This bar chart compares the Length (in kilobases, yellow bars) and Count (purple bars) of annotated features for specific regions across chromosomes in the genome. The x-axis represents distinct chromosome regions, including ChrB-R (Right arm of Chromosome B), ChrC-L (Left arm of Chromosome C), ChrC-R (Right arm of Chromosome C), ChrE-R (Right arm of Chromosome E), ChrF-L (Left arm of Chromosome F), ChrF-R (Right arm of Chromosome F). The yellow bars indicate the cumulative length of the regions (in kilobases), while the purple bars indicate the total count of rDNA repeats (excluding telomeric) identified within these regions. The rDNA repeats were determined by similarity to published rDNA sequences.

## DISCUSSION

Heterothallic yeast, like *Yarrowia lipolytica*, typically engage in outcrossing, where genetic material is exchanged between individuals of different mating types. This may lead to greater genetic diversity and adaptability to changing environments and contribute to divergence and speciation [61]. In contrast, homothallic yeast, such as *S. cerevisiae*, primarily engage in selfing since they can switch their mating type through a gene conversion process initiated by the HO endonuclease [62], where diploidization occurs within nearly identical genotypes. This process may result in the fixation of beneficial alleles or the accumulation of deleterious mutations, potentially leading to lower genetic diversity [63]. These differences in reproductive strategies lead to distinct recombination pathways in the two types of yeast [64]. However, a recent comparison of 56 shot-gun sequenced strains showed a very low level of genetic diversity, indicating that *Y. lipolytica* may be a species that has recently emerged [25].

*Y. lipolytica* exhibits a remarkably low rate of mating and spore viability between different lineages due to chromosomal rearrangements, which may contribute to its poor fertility. However, even without genome rearrangements, *Yarrowia* has very poor mating frequency [5]. Chromosomal rearrangements in *Y. lipolytica* could have been caused by crossing-over events facilitated by the different types of transposable elements present in the organism [27]. Yeast genomes contain mobile genetic elements, such as transposons and retrotransposons, which can translocate within the genome. These elements can be inserted into new locations within the genome or cause chromosomal reorganization by combining with various regions. The prevalence of repeated sequences found in the E122 *MATA* and 22301-5 *MATB* strains may also play a role in rearranging and evolving the genome of this yeast species, consistent with the notion that transposable elements and other repetitive elements can be significant contributors to genome evolution [25,27].

We found a higher number of introns in the E122 *MATA* and 22301-5 *MATB* strains compared to the DSM 3286 and CLIB122 strains (Figure 4), with most genes having a single intron. In particular, we observed a higher proportion of genes with more than one intron in E122 and 22301-5. This intron-rich genome may enable the production of multiple protein isoforms from a single gene, offering *Yarrowia* the ability to rapidly adapt to changing environments or industrial processes. This could prove especially valuable for *Yarrowia*, as it frequently operates in diverse and challenging growth conditions.

The enhanced genome assembly and annotation of *Yarrowia lipolytica* strain E122 *MATA* and 22301-5 *MATB*, was made possible by employing a hybrid sequencing approach that combined the precision of Illumina short reads with the length of Oxford Nanopore long reads and advanced base calling with Guppy 5. This high-quality assembly allowed us to capture telomeric regions, rDNA repeats and improve the completeness of essential gene markers, achieving a BUSCO score of 96.8%. These results establish a strong foundation for further functional and comparative studies on *Y. lipolytica* and its applications.

### Gene Annotation and Genetic Diversity

Our comparative gene analysis revealed significant differences between the sequenced E122 *MATA* and 22301-5 *MATB* strains and other previously studied *Yarrowia* strains, such as DSM 3286 and CLIB-122. The increased gene count in E122 *MATA* and 22301-5 *MATB* and the presence of alternative isoforms suggest a potentially broader genetic repertoire and greater regulatory complexity in these strains. This added complexity may reflect adaptive mechanisms developed in response to specific environmental and industrial conditions. For instance, the increased median gene length, attributed to a higher intron count, suggests unique gene structures that could enhance regulatory flexibility, supporting more intricate metabolic or stress-response pathways.

The core gene analysis indicates that E122 *MATA* and 22301-5 *MATB* share 5,975 core genes with other *Yarrowia* strains, consistent with prior findings of limited genetic diversity within *Yarrowia lipolytica* [25]. However, identifying 1,204 unique ortholog groups in *MATA* and *MATB* suggests subtle genomic differences that could contribute to strain-specific phenotypes. Gene Ontology (GO) enrichment analysis of these unique genes emphasizes metabolic processes and transcription initiation. These functions are advantageous in environmental settings where efficient resource utilization and adaptability to stressful conditions are beneficial.

### Repeat Elements and Genomic Evolution

The investigation into repetitive DNA elements highlights more repeats in E122 *MATA* and 22301-5 *MATB*, particularly in RNA, LTRs, LINE, and SINE elements, compared to DSM 3286 and CLIB122 (Figure 6B). The distinct repeat profiles observed in E122 *MATA* and 22301-5 *MATB*—with a balance of RNA and simple repeats making up 30-40% of the genome—suggest unique genomic architectures that may influence adaptation. The higher proportions of certain repeat types, particularly simple and RNA repeats, could facilitate rapid genomic changes, enhancing adaptability in dynamic environments like industrial fermentation.

These repeat variations among strains may indicate different genomic stability and plasticity strategies. For example, the high LTR content in DSM 3286 may signify historical transposon activity, promoting genomic rearrangements. In contrast, the more balanced and stable repeat landscape in E122 *MATA* and 22301-5 *MATB* suggests a refined evolutionary adaptation that could confer resilience in industrial contexts. The conserved nature of certain repeat types across strains, such as LINE elements, suggests shared functional roles across *Yarrowia* lineages, whereas the unique repeat profiles of E122 *MATA* and 22301-5 *MATB* reflect strain-specific evolutionary pressures.

### rDNA Repeats and Size

Analysis of the distribution of rDNA repeats in the E122 *MATA* strain revealed distinct patterns in repeat length and counts across multiple chromosome regions, adjacent to the telomeres as shown in Figure 7A. Regions chrC-R, chrE-R, and chrF-R exhibit the most extended total rDNA lengths (∼11 KB) with moderate counts, suggesting these regions contain larger rDNA repeats or a higher density of sizeable elements. In contrast, chrC-L has a shorter total length (∼8 KB) and a lower count, indicating fewer and potentially smaller rDNA repeats. This heterogeneity could indicate region-specific roles or stability requirements for rDNA within the MAT-A strain. We note that the location of the rDNA repeats in the two strains analyzed herein is essentially the same as in DSM 3286, but the estimated number of rDNA repeats differs. We suggest that this difference reflects both technical and biological variation.

The stability of the rDNA and telomeric repeats needs explanation. By stability we refer to the prevention of recombination between rDNA repeats that could reduce their number. In *S. cerevisiae*, the SIR proteins play an important role in the maintenance of the rDNA repeats by preventing recombination [14,65]. *Y. lipolytica* lacks the SIR proteins, except for *SIR2*, which is present in all eukaryotes, and it also lacks genes encoding RNAi components that in other eukaryotes suppress gene expression in heterochromatin [16,66]. This raises the interesting issue of how the rDNA and telomeric repeats resist recombination and thus maintain stability. One possibility is that SIR2 is sufficient for preventing recombination between repeats in *Y. lipolytica*, but we think this is unlikely since SIR2 is not sufficient in *S. cerevisiae*. Alternatively, the rDNA repeats are all adjacent to telomeric repeats in *Y. lipolytica* and it is possible that a telomere associated protein complex suppresses recombination in *cis*, well into the adjacent rDNA repeats.

### GC Content and Evolutionary Implications

The overall GC content in *Y. lipolytica* is higher than in *S. cerevisiae*, which aligns with the significant evolutionary divergence between these species, estimated at around 300 million years. The consistent GC content across E122 *MATA*, 22301-5 *MATB*, and DSM 3286 (48.9%) compared to the low GC content of the mitochondrial chromosome (22.59%) suggests differences in selective pressures and genome organization between nuclear and mitochondrial genomes. This higher GC content may have implications for DNA stability, transcription efficiency, and DNA replication dynamics, offering insights into the evolutionary and functional constraints on the *Yarrowia* genome. For example, the well-characterized origins of DNA replication in *S. cerevisiae* are AT-rich. We are analyzing the genome replication and origins of DNA replication in *Yarrowia* to determine if the genome has GC-rich origins of DNA replication that are more akin to the GC-rich origins in human cells. The more complete genome sequences of the E122 *MATA* and 22301-5 *MATB* strains should facilitate the analysis of genome replication patterns and mechanisms.

### Conclusions and Future Directions

Our findings provide a comprehensive view of the genomic landscape and diversity within *Yarrowia lipolytica* strains *MATA* and *MATB*, laying the groundwork for further research in functional genomics and strain optimization. The variability in repeat elements, the distinct genomic organization, and the elevated gene complexity observed in *MATA* and *MATB* highlight the evolutionary and functional divergence within *Y. lipolytica*. The insights gained from future studies of basic molecular biology in *Y. lipolytica* will contribute to our understanding of the molecular underpinnings that enable *Yarrowia lipolytica* to thrive in highly varied environments, ultimately advancing strain development for biotechnology. The additional genes identified in this genome analysis may provide new metabolic functions that will facilitate engineering of new chemical pathways for industrial use.

## DATA AVAILABILITY

The DNA sequence and annotation data are available at Dryad

https://doi.org/10.5061/dryad.v15dv427f.The data are:

***Files and variables***

**File: yarrowia_lipolytica-22301-5.fasta Description:** Strain 22301-5 Genome assembly

**File: yarrowia_lipolytica-E122.fasta Description:** Strain E122 Genome assembly

**File: yarrowia_lipolytica-E122.gff**

Description: Strain E122 Transcriptome assembly and annotation. Full structural gene annotation in GFF3 format (genes, exons, CDS)

**File: yarrowia_lipolytica-22301-5.gff**

Description: Strain 22301-5 Transcriptome assembly and annotation. Full structural gene annotation in GFF3 format (genes, exons, CDS)

**File: yarrowia_lipolytica-22301-5.AED_filter_1.gff**

Description: Strain 22301-5 Filtered GFF with high-confidence gene models (AED ≤ 1).

**File: yarrowia_lipolytica-E122.AED_filter_1.gff**

Description: Strain E122 Filtered GFF with high-confidence gene models (AED ≤ 1).

**File: yarrowia_lipolytica-22301-5.evd.cdna.fasta**

Description: Strain 22301-5 FASTA of spliced cDNA (transcripts including UTRs).

**File: yarrowia_lipolytica-E122.evd.cdna.fasta**

Description: Strain E122 FASTA of spliced cDNA (transcripts including UTRs).

**File: yarrowia_lipolytica-22301-5.evd.cds.fasta**

Description: Strain 22301-5 FASTA of coding sequences (CDS only).

**File: yarrowia_lipolytica-E122.evd.cds.fasta**

Description: Strain E122 FASTA of coding sequences (CDS only).

**File: yarrowia_lipolytica-22301-5.evd.protein.fasta**

Description: Strain 22301-5 FASTA of translated protein sequences.

**File: yarrowia_lipolytica-E122.evd.protein.fasta**

Description: Strain E122 FASTA of translated protein sequences.

**File: yarrowia_lipolytica-22301-5.evd.protein.interproscan.tsv**

Description: Strain 22301-5 InterProScan results: protein domain/function annotations.

**File: yarrowia_lipolytica-E122.evd.protein.interproscan.tsv**

Description: Strain E122 InterProScan results: protein domain/function annotations.

**File: yarrowia_lipolytica-22301-5.fasta.mod.EDTA.intact.gff3**

Description: Strain 22301-5 GFF3 of intact transposable elements (TEs) identified by EDTA.

**File: yarrowia_lipolytica-E122.fasta.mod.EDTA.intact.gff3**

Description: Strain E122 GFF3 of intact transposable elements (TEs) identified by EDTA.

**File: yarrowia_lipolytica-22301-5.fasta.mod.EDTA.TEanno.gff3**

Description: Strain 22301-5 GFF3 of all annotated TE features (both intact and fragmented).

**File: yarrowia_lipolytica-E122.fasta.mod.EDTA.TEanno.gff3**

Description: Strain E122 GFF3 of all annotated TE features (both intact and fragmented).

**File: yarrowia_lipolytica-22301-5.fasta.mod.EDTA.TEanno.sum**

Description: Strain 22301-5 Summary statistics of TE annotations (counts, lengths, classes).

**File: yarrowia_lipolytica-E122.fasta.mod.EDTA.TEanno.sum**

Description: Strain E122 Summary statistics of TE annotations (counts, lengths, classes).

**File: yarrowia_lipolytica-22301-5.fasta.mod.EDTA.TElib.fa**

Description: Strain 22301-5 Custom TE library (consensus sequences of identified repeats).

**File: yarrowia_lipolytica-E122.fasta.mod.EDTA.TElib.fa**

Description: Strain E122 Custom TE library (consensus sequences of identified repeats).

**File: yarrowia_lipolytica-22301-5.fasta.mod.MAKER.masked**

Description: Strain 22301-5 Genome sequence with repeats masked (soft-masked by MAKER using EDTA TE annotations).

**File: yarrowia_lipolytica-E122.fasta.mod.MAKER.masked**

Description: Strain E122 Genome sequence with repeats masked (soft-masked by MAKER using EDTA TE annotations).

**File: repeatLib.fungi.fa**

Description: Custom fungal repeat library in FASTA format (used for repeat masking).

**File: yarrowia_lipolytica-22301-5_core_2_87_1.repeat_report**

Description: Strain 22301-5 RepeatMasker output report summarizing repeat content (type, count, coverage).

**File: yarrowia_lipolytica-E122_core_2_87_1.repeat_report**

Description: Strain E122 RepeatMasker output report summarizing repeat content (type, count, coverage).

**File: yarrowia_lipolytica-22301-5_repeat_feature.gene.overlap.txt** Description: Strain 22301-5 TE-related genes overlapping repeat element

**File: yarrowia_lipolytica-E122_repeat_feature.gene.overlap.txt** Description: Strain E122 TE-related genes overlapping repeat element.

**File: yali_id.system_name.orthologs.csv**

Description: CSV file containing ortholog relationships between *Yarrowia lipolytica* genes and genes from other species or strains; includes gene IDs and matched orthologs.

**Code/software**

Genome and transcriptome data, as well as gff3 annotation files can all be viewed using a genome browser. (e.g. IGV). The remaining files can be opened in a text editor/viewer or excel.

## AUTHOR CONTRIBUTIONS

N.Z. did the experiments and the DNA sequencing using the Cold Spring Harbor Laboratory Sequencing and Technology Core facility. O. E D., K.C., Z.L. and D.W. performed data analysis. B.S. conceived the project and oversaw all aspects of the research. N.Z. K.C. D.W. and B.S. wrote the paper.

## ACKNOWLEDGEMENTS

The authors thank Dr. Richard A. Rachubunski, University of Alberta, Canada for providing the *Yarrowia* strains used in this project. We thank members of the Cold Spring Harbor Laboratory Sequencing and Technology Core facility for DNA sequence analysis.

## FUNDING

Funding of this research was supported by a grant (GM45436) to B.S. from the National Institute of General Medical Sciences at the National Institutes of Health and a grant (8062-21000-051-000D) to D.W. from the United States Department of Agriculture (USDA), Agricultural Research Service (ARS). USDA ARS. The Cold Spring Harbor Laboratory Genome Sequencing and Technologies Core was supported in part by the Cold Spring Harbor Laboratory Cancer Center grant (CA13106) from the National Cancer Institute at the National Institutes of Health.

## CONFLICT OF INTEREST

The authors declare no conflict of interest.

## Notes

https://doi.org/10.5061/dryad.v15dv427f

### Competing Interest Statement

The authors have declared no competing interest.

### Summary of Updates

This version of the paper has been updated with new analysis of the two Yarrowia genomes and public release of the genome data.

## REFERENCES

1. Gonçalves FAG, Colen G, Takahashi JA. Yarrowia lipolytica and Its Multiple Applications in the Biotechnological Industry. Sci World J. 2014;2014(1):476207.

2. Mamaev D, Zvyagilskaya R. Yarrowia lipolytica : A multitalented yeast species of ecological significance. Fems Yeast Res. 2021;

3. Bankar AV, Kumar AR, Zinjarde SS. Environmental and industrial applications of Yarrowia lipolytica. Appl Microbiol Biotechnol. 2009;84(5):847.

4. Groenewald M, Boekhout T, Neuvéglise C, Gaillardin C, Dijck PWM van, Wyss M. Yarrowia lipolytica: Safety assessment of an oleaginous yeast with a great industrial potential. Crit Rev Microbiol. 2014;40(3):187– 206.

5. Barth G, Gaillardin C. Physiology and genetics of the dimorphic fungus Yarrowia lipolytica. Fems Microbiol Rev. 1997 Apr;19(4):219–37.

6. Magnan C, Yu J, Chang I, Jahn E, Kanomata Y, Wu J, Zeller M, Oakes M, Baldi P, Sandmeyer S. Sequence Assembly of Yarrowia lipolytica Strain W29/CLIB89 Shows Transposable Element Diversity. Plos One. 2016;11(9):e0162363.

7. Fumasoni M, Murray AW. The evolutionary plasticity of chromosome metabolism allows adaptation to constitutive DNA replication stress. Elife. 2020 Feb 11;9:e51963.

8. Mekouar M, Blanc-Lenfle I, Ozanne C, Silva CD, Cruaud C, Wincker P, Gaillardin C, Neuvéglise C. Detection and analysis of alternative splicing in Yarrowia lipolytica reveal structural constraints facilitating nonsense-mediated decay of intron-retaining transcripts. Genome Biol. 2010;11(6):R65.

9. Hu Y, Tareen A, Sheu YJ, Ireland WT, Speck C, Li H, Joshua-Tor L, Kinney JB, Stillman B. Evolution of DNA replication origin specification and gene silencing mechanisms. Nat Commun. 2020;11(1):5175.

10. Marahrens Y, Stillman B. A yeast chromosomal origin of DNA replication defined by multiple functional elements. Science (New York, NY) [Internet]. 1992 Feb 14;255(5046):817–23. Available from: http://www.sciencemag.org/cgi/reprint/255/5046/817

11. Bell SP, Labib K. Chromosome Duplication in Saccharomyces cerevisiae. Genetics. 2016 July 6;203(3):1027–67.

12. Lee CSK, Cheung MF, Li J, Zhao Y, Lam WH, Ho V, Rohs R, Zhai Y, Leung D, Tye BK. Humanizing the yeast origin recognition complex. Nat Commun. 2021;12(1):33.

13. Rusche, L. N. and Hickman, M. A. Evolution of Silencing at the Mating-type loci in Hemiascomycetes, In Sex in Fungi: Molecular Determination and Evolutionary Implications. pp. 189–200. Edited by Joseph Heitman, James W. Kronstad, John W. Taylor, Lorna A. Casselton. 2007 ASM Press, Washington, D.C.

14. Rusche LN, Kirchmaier AL, Rine J. The establishment, inheritance, and function of silenced chromatin in Saccharomyces cerevisiae. Annual review of biochemistry. 2003;72(1):481–516.

15. Hickman MA, Froyd CA, Rusche LN. Reinventing heterochromatin in budding yeasts: Sir2 and the origin recognition complex take center stage. Eukaryotic cell. 2011 Sept;10(9):1183–92.

16. Smith JS, Brachmann CB, Celic I, Kenna MA, Muhammad S, Starai VJ, Avalos JL, Escalante-Semerena JC, Grubmeyer C, Wolberger C, Boeke JD. A phylogenetically conserved NAD+-dependent protein deacetylase activity in the Sir2 protein family. Proc Natl Acad Sci. 2000;97(12):6658–63.

17. Bell SP, Kobayashi R, Stillman B. Yeast Origin Recognition Complex Functions in Transcription Silencing and DNA Replication. Science. 1993;262(5141):1844–9.

18. Hickman MA, Rusche LN. Transcriptional silencing functions of the yeast protein Orc1/Sir3 subfunctionalized after gene duplication. Proceedings of the National Academy of Sciences. 2010 Nov 9;107(45):19384–9.

19. Hou Z, Bernstein DA, Fox CA, Keck JL. Structural basis of the Sir1–origin recognition complex interaction in transcriptional silencing. Proc Natl Acad Sci. 2005;102(24):8489–94.

20. Grunstein M, Gasser SM. Epigenetics in Saccharomyces cerevisiae. Cold Spring Harbor Perspectives in Biology. 2013 July 1;5(7):a017491.

21. Maria H, Rusche LN. The DNA replication protein Orc1 from the yeast Torulaspora delbrueckii is required for heterochromatin formation but not as a silencer-binding protein. Genetics. 2022;222(1):iyac110.

22. Hu Y, Stillman B. Origins of DNA replication in eukaryotes. Mol Cell. 2023;83(3):352–72.

23. Hyrien O, Guilbaud G, Krude T. The double life of mammalian DNA replication origins. Genes Dev. 2025;39:304–24.

24. Devillers H, Neuvéglise C. Genome Sequence of the Oleaginous Yeast Yarrowia lipolytica H222. Microbiol Resour Announc. 2019;8(4):10.1128/mra.01547-18.

25. Bigey F, Pasteur E, Połomska X, Thomas S, Coq AMCL, Devillers H, Neuvéglise C. Insights into the Genomic and Phenotypic Landscape of the Oleaginous Yeast Yarrowia lipolytica. J Fungi. 2023;9(1):76.

26. Liu L, Alper HS. Draft Genome Sequence of the Oleaginous Yeast Yarrowia lipolytica PO1f, a Commonly Used Metabolic Engineering Host. Genome Announc. 2014;2(4):e00652–14.

27. Luttermann T, Rückert C, Wibberg D, Busche T, Schwarzhans JP, Friehs K, Kalinowski J. Establishment of a near-contiguous genome sequence of the citric acid producing yeast Yarrowia lipolytica DSM 3286 with resolution of rDNA clusters and telomeres. Nar Genom Bioinform. 2021;3(4):lqab085-.

28. Devillers H, Brunel F, Połomska X, Sarilar V, Lazar Z, Robak M, Neuvéglise C. Draft Genome Sequence of Yarrowia lipolytica Strain A-101 Isolated from Polluted Soil in Poland. Genome Announc. 2016;4(5):e01094–16.

29. Madzak C. Yarrowia lipolytica Strains and Their Biotechnological Applications: How Natural Biodiversity and Metabolic Engineering Could Contribute to Cell Factories Improvement. J Fungi. 2021;7(7):548.

30. Barth G, Gaillardin C. Yarrowia Lipolytica. In Nonconventional Yeasts in Biotechnology, A Handbook. In: Wolf K, editor. Springer Berlin, Heidelberg, New York; 1996. p. 313–88.

31. Mansour S, Bailly J, Landaud S, Monnet C, Sarthou AS, Cocaign-Bousquet M, Leroy S, Irlinger F, Bonnarme P. Investigation of Associations of Yarrowia lipolytica, Staphylococcus xylosus, and Lactococcus lactis in Culture as a First Step in Microbial Interaction Analysis. Appl Environ Microbiol. 2009;75(20):6422–30.

32. Kolmogorov M, Yuan J, Lin Y, Pevzner PA. Assembly of long, error-prone reads using repeat graphs. Nat Biotechnol. 2019;37(5):540–6.

33. Martin, M. Cutadapt removes adapter sequences from high-throughput sequencing reads. EMBnet.journal [SI] [Internet]. 2011;(17):10–2. Available from: https://journal.embnet.org/index.php/embnetjournal/article/view/200/479

34. Li H, Durbin R. Fast and accurate short read alignment with Burrows–Wheeler transform. Bioinformatics. 2009;25(14):1754–60.

35. Danecek P, Bonfield JK, Liddle J, Marshall J, Ohan V, Pollard MO, Whitwham A, Keane T, McCarthy SA, Davies RM, Li H. Twelve years of SAMtools and BCFtools. GigaScience. 2021;10(2):giab008.

36. Walker BJ, Abeel T, Shea T, Priest M, Abouelliel A, Sakthikumar S, Cuomo CA, Zeng Q, Wortman J, Young SK, Earl AM. Pilon: An Integrated Tool for Comprehensive Microbial Variant Detection and Genome Assembly Improvement. PLoS ONE. 2014;9(11):e112963.

37. Alonge M, Soyk S, Ramakrishnan S, Wang X, Goodwin S, Sedlazeck FJ, Lippman ZB, Schatz MC. RaGOO: fast and accurate reference-guided scaffolding of draft genomes. Genome Biol. 2019;20(1):224.

38. Bolger AM, Lohse M, Usadel B. Trimmomatic: a flexible trimmer for Illumina sequence data. Bioinformatics. 2014;30(15):2114–20.

39. Pertea M, Pertea GM, Antonescu CM, Chang TC, Mendell JT, Salzberg SL. StringTie enables improved reconstruction of a transcriptome from RNA-seq reads. Nat Biotechnol. 2015;33(3):290–5.

40. Kainth AS, Haddad GA, Hall JM, Ruthenburg AJ. Merging short and stranded long reads improves transcript assembly. PLOS Comput Biol. 2023;19(10):e1011576.

41. Pertea G, Pertea M. GFF Utilities: GffRead and GffCompare. F1000Research. 2020;9:ISCB Comm J-304.

42. Campbell MS, Holt C, Moore B, Yandell M. Genome Annotation and Curation Using MAKER and MAKER-P. Curr Protoc Bioinform. 2014;48(1):4.11.1–4.11.39.

43. Huang Y, Niu B, Gao Y, Fu L, Li W. CD-HIT Suite: a web server for clustering and comparing biological sequences. Bioinformatics. 2010;26(5):680–2.

44. Trincado JL, Entizne JC, Hysenaj G, Singh B, Skalic M, Elliott DJ, Eyras E. SUPPA2: fast, accurate, and uncertainty-aware differential splicing analysis across multiple conditions. Genome Biol. 2018;19(1):40.

45. Solovyev V, Kosarev P, Seledsov I, Vorobyev D. Automatic annotation of eukaryotic genes, pseudogenes and promoters. Genome Biol. 2006;7(Suppl 1):S10.

46. Stanke M, Diekhans M, Baertsch R, Haussler D. Using native and syntenically mapped cDNA alignments to improve de novo gene finding. Bioinformatics. 2008;24(5):637–44.

47. Ou S, Su W, Liao Y, Chougule K, Agda JRA, Hellinga AJ, Lugo CSB, Elliott TA, Ware D, Peterson T, Jiang N, Hirsch CN, Hufford MB. Benchmarking transposable element annotation methods for creation of a streamlined, comprehensive pipeline. Genome Biol. 2019;20(1):275.

48. Haas BJ, Delcher AL, Mount SM, Wortman JR, Jr RKS, Hannick LI, Maiti R, Ronning CM, Rusch DB, Town CD, Salzberg SL, White O. Improving the Arabidopsis genome annotation using maximal transcript alignment assemblies. Nucleic Acids Res. 2003;31(19):5654–66.

49. Jones P, Binns D, Chang HY, Fraser M, Li W, McAnulla C, McWilliam H, Maslen J, Mitchell A, Nuka G, Pesseat S, Quinn AF, Sangrador-Vegas A, Scheremetjew M, Yong SY, Lopez R, Hunter S. InterProScan 5: genome-scale protein function classification. Bioinformatics. 2014;30(9):1236–40.

50. Olson AJ, Ware D. Ranked choice voting for representative transcripts with TRaCE. Bioinformatics. 2021;38(1):261–4.

51. Stabenau A, McVicker G, Melsopp C, Proctor G, Clamp M, Birney E. The Ensembl Core Software Libraries. Genome Res. 2004;14(5):929–33.

52. Vilella AJ, Severin J, Ureta-Vidal A, Heng L, Durbin R, Birney E. EnsemblCompara GeneTrees: Complete, duplication-aware phylogenetic trees in vertebrates. Genome Res. 2009;19(2):327–35.

53. Simão FA, Waterhouse RM, Ioannidis P, Kriventseva EV, Zdobnov EM. BUSCO: assessing genome assembly and annotation completeness with single-copy orthologs. Bioinformatics. 2015;31(19):3210–2.

54. Pérez-Wohlfeil E, Diaz-del-Pino S, Trelles O. Ultra-fast genome comparison for large-scale genomic experiments. Sci Rep. 2019;9(1):10274.

55. Rosas-Quijano R, Gaillardin C, Ruiz-Herrera J. Functional Analysis of the MATB Mating-Type Idiomorph of the Dimorphic Fungus Yarrowia lipolytica. Curr Microbiol. 2008 Aug;57(2):115–20.

56. Kurischko C, Schilhabel MB, Kunze I, Franzl E. The MAT A locus of the dimorphic yeast Yarrowia lipolytica consists of two divergently oriented genes. Mol Gen Genet MGG. 1999;262(1):180–8.

57. Nuttley WM, Brade AM, Eitzen GA, Glover JR, Aitchison JD, Gaillardin C. Rapid identification and characterization of peroxisomal assembly mutants in Yarrowia lipolytica. Yeast (Chichester, England). 1993 May 1;9(5):507–17.

58. Dujon B, Sherman D, Fischer G, Durrens P, Casaregola S, Lafontaine I, Montigny J de, Marck C, Neuvéglise C, Talla E, Goffard N, Frangeul L, Aigle M, Anthouard V, Babour A, Barbe V, Barnay S, Blanchin S, Beckerich JM, Beyne E, Bleykasten C, Boisramé A, Boyer J, Cattolico L, Confanioleri F, Daruvar A de, Despons L, Fabre E, Fairhead C, Ferry-Dumazet H, Groppi A, Hantraye F, Hennequin C, Jauniaux N, Joyet P, Kachouri R, Kerrest A, Koszul R, Lemaire M, Lesur I, Ma L, Muller H, Nicaud JM, Nikolski M, Oztas S, Ozier-Kalogeropoulos O, Pellenz S, Potier S, Richard GF, Straub ML, Suleau A, Swennen D, Tekaia F, Wésolowski-Louvel M, Westhof E, Wirth B, Zeniou-Meyer M, Zivanovic I, Bolotin-Fukuhara M, Thierry A, Bouchier C, Caudron B, Scarpelli C, Gaillardin C, Weissenbach J, Wincker P, Souciet JL. Genome evolution in yeasts. Nature. 2004;430(6995):35–44.

59. Peona V, Blom MPK, Xu L, Burri R, Sullivan S, Bunikis I, Liachko I, Haryoko T, Jønsson KA, Zhou Q, Irestedt M, Suh A. Identifying the causes and consequences of assembly gaps using a multiplatform genome assembly of a bird-of-paradise. Mol Ecol Resour. 2021;21(1):263–86.

60. Baptista RP, Reis-Cunha JL, DeBarry JD, Chiari E, Kissinger JC, Bartholomeu DC, Macedo AM. Assembly of highly repetitive genomes using short reads: the genome of discrete typing unit III Trypanosoma cruzi strain 231. Microb Genom. 2018;4(4):e000156.

61. Lee SC, Ni M, Li W, Shertz C, Heitman J. The Evolution of Sex: a Perspective from the Fungal Kingdom. Microbiol Mol Biol Rev. 2010;74(2):298–340.

62. Haber JE. Mating-Type Genes and MAT Switching in Saccharomyces cerevisiae. Genetics. 2012;191(1):33– 64.

63. Hanson SJ, Wolfe KH. An Evolutionary Perspective on Yeast Mating-Type Switching. Genetics. 2017 May;206(1):9–32.

64. Dujon BA, Louis EJ. Genome Diversity and Evolution in the Budding Yeasts (Saccharomycotina). Genetics. 2017 June;206(2):717–50.

65. Kueng S, Oppikofer M, Gasser SM. SIR proteins and the assembly of silent chromatin in budding yeast. Annual review of genetics. 2013;47:275–306.

66. Blander G, Guarente L. The SIR2 Family of Protein Deacetylases. Biochemistry. 2004;73(1):417–35.

